# Innate Defense Mechanisms Against *Nosema ceranae* in Hygienic Honey Bee (*Apis mellifera*) Colonies

**DOI:** 10.64898/2026.02.02.693565

**Authors:** M. Sydney Miller, Dawn Boncristiani, Jay Evans, P. Alexander Burnham, Cailin Barrett, Kaira Wagoner, Samantha A. Alger

## Abstract

The honey bee colony (*Apis mellifera*) acts as a superorganism, with a dual immune system that operates at the individual and social level. However, the linkages between immune mechanisms across the two levels remain poorly understood, despite the relevance for developing effective breeding strategies to improve honey bee disease resistance. Hygienic behavior involving the removal of unhealthy brood is a key component of honey bee social immunity and is highly effective at limiting parasites and pathogens in the colony. While this form of hygienic behavior can reduce brood diseases, parasites infecting adult bees primarily, such as *Nosema ceranae,* are not directly impacted by the behavior. However, when using the Unhealthy Brood Odor (UBeeO) assay to quantify hygienic behavior performance, hygienic colonies have been shown to maintain lower *Nosema* spp. loads over time and overall compared to non-hygienic colonies. To investigate the mechanisms driving reduced *Nosema* spp. in hygienic colonies, we conducted a series of field and lab experiments to test the innate immune performance of individual bees. We evaluated several factors across hygienic and non-hygienic bees including (1) differences in *N. ceranae* infection levels, (2) survival probability, (3) *Vitellogenin* and *Hymenoptaecin* gene expression, and (4) amount of *N. ceranae* inoculant consumed. We found that hygienic bees consumed less of the inoculant, exhibited upregulated *Vitellogenin* gene expression at peak *N. ceranae* infection, showed a positive relationship between *Hymenoptaecin* gene expression and *N. ceranae* infection levels, and had greater survivability when infected with *N. ceranae*, compared to non-hygienic bees. Here, we present new findings that link colony hygienic behavior performance to individual-level resistance and tolerance mechanisms in response to *N. ceranae*, suggesting broader implications for the success of selective breeding programs targeting hygienic traits.

## Introduction

Pests and pathogens are a primary threat to honey bees (*Apis mellifera*) impacting the health of brood and adult bees and contributing to overall colony decline. In response to intruders, the honey bee colony acts as a superorganism, with a dual immune system that operates at the individual and social level [1–3]. Honey bees rely on their innate immune system (e.g. physical barriers, cellular and humoral immunity) to defend against infection as well as complex social behaviors that reduce the impacts of parasites and pathogens in colonies. Understanding the linkages between immune function at the social and individual level is essential for informing effective selective breeding strategies aimed at improving honey bee disease resistance and colony survival.

Hygienic behavior refers to the enhanced ability of worker bees to respond to chemical odorants emitted by diseased or dead brood (developing larvae and/or pupae) by uncapping and removing pupae from the nest [4,5]. The form of hygienic behavior involving the removal of unhealthy brood from the hive should be distinguished from other forms of honey bee hygiene, such as known auto- and allo-grooming behaviors performed by adults [1,6]. As a heritable genetic trait, hygienic behavior is among the most important social behaviors for conferring colony-level resistance against brood diseases [7–9] and in recent years has become a major focus in honey bee breeding programs. Previously developed assays used to quantify hygienic behavior (e.g. pin prick, freeze-killed brood) are based in necrophoric activity and have shown to confer reduced levels of Foulbrood, Chalkbrood, and *Varroa* infestations [10–13]. As an improved method for quantifying hygienic behavior, the Unhealthy Brood Odor (UBeeO) assay challenges bees with synthetic pheromones mimicking the natural odors emitted by live, parasitized brood. In addition to predicting a low incidence of brood disease, the UBeeO assay has been shown to predict lower spore loads of *Nosema* spp. over time and overall, in hygienic colonies compared to non-hygienic colonies [14].

*Nosema ceranae* is a common microsporidian endoparasite that infects the midgut epithelial cells [15,16] of adult bees [17,18]. *Nosema ceranae* infection negatively impacts honey bee health at the individual level—causing nutritional and energetic stress [19,20], immunosuppression [21,22], altered behavior [23,24], reduced lifespan, and inhibition of host cell apoptosis [25–27]—which can reduce colony fitness by lowering brood numbers and honey production, and in severe cases, lead to colony death [15,28]. With limited viable treatment options available to beekeepers, effective prevention and colony management remain essential for controlling the pathogen [28]. Moreover, the risk of target pests and pathogens building resistance to chemicals and rendering treatments ineffective—as seen in global *Varroa* populations resistant to several well-known acaricides [29–31]—further underscores the need for more sustainable interventions to control honey bee pests and diseases.

*Nosema ceranae* is primarily an adult bee pathogen [32,33] that has only been found to infect brood through manual inoculation in lab studies [34,35] or at extremely low prevalences (1-3%) in natural hive settings [36,37]. While *N. ceranae* infection in developing brood has not been thoroughly evaluated, many studies have reported an absence of *N. ceranae* infection in emerging adults [17,38,39], suggesting that brood does not normally become infected in the hive. Since hygienic behavior acts on infected brood, it has not been shown to directly inhibit *Nosema* spp. transmission, aside from reducing its prevalence at the apiary level [14,40]. In a hygiene-based breeding program in Turkey [40], researchers reported that average apiary-level hygienic behavior increased from 43% to 93% (n = 123), while *Nosema* spp. levels declined consistently from 61% to 19% in only three years. Therefore, recent findings by Alger et al. [14] are not the first to demonstrate an association between high hygienic behavior and low *Nosema* spp. incidence in a field study. Nevertheless, it remains unclear how hygienic colonies maintain low *Nosema* spp. loads and whether colony-level resistance arises from social immunity in the form of pleiotropic effects on brood and adult bee hygiene, innate immune mechanisms, or a combination of both.

Several co-occurring mechanisms at the individual and social level may contribute to colony-level resistance to *N. ceranae*. At the social level, adults in hygienic colonies may communicate their diseased state through stronger chemical signals, prompting detection and removal by nestmates, similar to the process of removing brood [7,41]. Adults in hygienic colonies may, in turn, be more sensitive to atypical odors and better able to detect, isolate, and/or discard of *N. ceranae*-contaminated individuals and/or food sources in the hive. To better understand the social dynamics of *N. ceranae*-infected bees in hygienic colonies, it is necessary to first evaluate their innate performance against *N. ceranae*.

Previous studies have shown no genetic tradeoffs between hygienic behavior and innate immunity [42]. In fact, hygiene-performing bees have been associated with modifications to the gut microbiome [43] and higher expression of antimicrobial peptides [44], which may lead to more efficient immune responses against invading pathogens. Therefore, individuals in hygienic colonies may exhibit stronger innate immunity and improved performance under pathogen stress overall. Specifically, they might lessen the negative health impacts of *N. ceranae* infection by upregulating immune-related genes in cellular or humoral pathways, which could help limit pathogen invasion or tissue damage [12,42].

Two major immune genes that are likely responsible for enhancing innate performance against *N. ceranae* are *Hymenoptaecin* and *Vitellogenin*. *Hymenoptaecin* (Hym) is an antimicrobial peptide activated by the humoral immune system (*Imd* pathway) that directly resists pathogens by attacking their cell membranes [45]. *Vitellogenin* (Vg) is an egg yolk precursor protein that can repair tissue damage [46,47] and perform immunological defense functions against pathogens and reactive oxygen species [48,49]. *Vitellogenin* also influences multiple physiological functions in honey bees including behavioral maturation [50], social organization [51], longevity [52], and egg development [49]. Both Hym [22,53] and Vg are commonly downregulated in honey bees infected with *N. ceranae* or other parasites [54,55]; therefore, may play a central role in resisting *N. ceranae* infection in bees from hygienic colonies.

In this study, we investigated innate immune mechanisms that may enhance individual performance against *N. ceranae* infection and help explain the reduced *N. ceranae* loads observed in hygienic colonies in previous field studies. We compared bees from hygienic and non-hygienic colonies by evaluating (1) *N. ceranae* infection levels, (2) survival probability, (3) *Vitellogenin* and *Hymenoptaecin* gene expression, and (4) amount of *N. ceranae* inoculant consumed. Our objective was to identify individual-level traits associated with bees from hygienic colonies, which could suggest broad-spectrum disease management potential in hygiene-based breeding scenarios.

## Methods

### Pupal evaluations

Pupal samples were collected from *Nosema* spp.–infected honey bee colonies in St. Albans, Vermont and analyzed for spore presence to determine whether developing pupae experience infection under natural hive conditions. Since hygienic behavior targets unhealthy brood, it was important to determine whether pupae serve as a source of *Nosema* spp. infection in our target honey bee population and whether removing infected brood could help reduce pathogen loads in the colonies. Thirty pupae were collected in composite samples from each of 28 colonies with detectable *Nosema* spp. loads in nurse bees ranging from 5×10^4^ to 1.4×10⁶ spores per bee. Pink to purple-eyed pupae were extracted from their wax cell with forceps and stored at -80°C until processing [36].

To conduct spore counts on pupae, composite pupal samples were rinsed in phosphate buffered saline and pulverized in a plastic bag using a rolling pin. One mL of distilled water per pupa was added and allowed to settle for 45 s. Ten µL was transferred from the stock solution onto two haemocytometer (Improved Neubauer) counting chambers. Spores were counted under 40× magnification and converted to spores per pupae [56].

### Incubation and *N. ceranae* isolation

To compare hygienic and non-hygienic bees in our innate immune-response trials, we obtained newly emerged adults from hygienic and non-hygienic colonies. Throughout the text, we use the terms *hygienic bees* and *non-hygienic bees* to refer to individuals originating from hygienic and non-hygienic colonies, respectively, rather than bees actively performing hygienic behaviors. We constructed frame cages to house deep hive frames of emerging brood for 1-3 days. The frame cages consisted of a wooden frame and lid with 8 mm mesh screened sides that provided adequate ventilation. Adult bees were transferred to hoarding cages upon emergence (within ∼6 hours) to avoid consumption of contaminated food stores from their frames. We constructed hoarding cages to house adult workers for 12-14 days during *N. ceranae* spore inoculation and infection period. Each hoarding cage consisted of a 473 mL plastic cup with ventilation holes encircling the upper and lower rim. Feeders consisted of 5 mL plastic pipettes severed at the base of the bulb and secured in the straw hole of the plastic cup lid. A small piece of wax foundation served as a ramp to the feeder [56].

Adult workers in hoarding cages were maintained in two separate incubators to segregate *N. ceranae-*infected bees from non-infected control bees, both in complete darkness at 30°C and approximately 60-70% RH. A thermometer/humidity gauge was used to monitor the interior environmental conditions each day. Adult workers were fed a diet of 50% (v/v) sugar syrup that was administered via a pipette feeder. Fresh sugar syrup was replaced every other day during the infection period [56].

To obtain active *N. ceranae* spores for inoculation, we collected foragers [38] from live colonies with existing *Nosema* spp. loads of 11-13 million spores per bee. Foragers were collected in a package cage, fed 50% (v/v) sugar syrup, and placed in an incubator until processed. *Nosema ceranae* spores were isolated from 100 forager bees by homogenizing one bee per 0.5-1 mL of water. The solution was strained through 70 μm mesh, evaluated for concentration using standard microscopy and hemocytometer (Improved Neubauer), and diluted into 50% (v/v) sugar syrup to achieve spore concentrations of 10^4^ spores per 0.04 mL (low dose) or 5 × 10^4^ spores per 0.04 mL (high dose). Final inoculants were fed to bees the same day. Control bees received pure 50% (v/v) sugar syrup.

### Determining *Nosema* spp. inoculation method

To determine the most effective *Nosema* spp. inoculation strategy for our individual immune-response trials, we conducted a pilot study examining how *Nosema* spp. load and its variability are influenced by (1) individual versus group feeding and (2) the number of bees per cage under group-feeding conditions. Newly emerged adult bees were randomly assigned to one of the two feeding methods and, if assigned to group-feeding, cages of 30 or 10 bees. All bees were starved for 2-4 hours before administering the *Nosema* spp. inoculant. The *Nosema* spp. inoculant was administered to group-fed bees via ∼3 mL of sugar syrup containing 5 × 10^4^ spores per 0.04 mL, *ad libitum*, for 24 hours [57]. We used a pipette to administer 5 µL of sugar syrup containing 5 × 10^4^ spores to individually-fed bees, then returned the bees to their hoarding cages after 30 minutes (10 bees per cage) [33]. All bees were expected to consume 5 × 10^4^ spores by the end of the inoculation period. Bees were maintained in their hoarding cages for a 10-day infection period, at which time, bees were extracted from their cages and stored individually at -20°C until processed.

To conduct spore counts on adult bees using standard microscopy, the abdomens of individual bees were dissected, rinsed in phosphate buffered saline, and pulverized using a 1.5mL pestle for 90 s. One mL of distilled water was added and allowed to settle for 45 s. Ten µL was transferred from the stock solution onto two haemocytometer (Improved Neubauer) counting chambers. Spores were counted under 40× magnification and converted to spores per bee [56].

### Unhealthy Brood Odor (UBeeO) assays

To identify hygienic and non-hygienic colonies from which to source newly emerged adults for our innate immune-response trials, we tested 30 honey bee colonies located in Northern Vermont, which were part of an existing three-year program designed to select for hygienic behavior. Queens were reared and overwintered in Vermont and tested prior to the experiment in early June. The queens were primarily Carniolan (*Apis mellifera carnica*) and were not sourced from a designated “hygienic” line. No official permits were required to conduct hygienic behavior testing or pathogen sampling on live colonies, other than permission from Michael Palmer of French Hill Apiaries and Bianca Braman of Vermont Bees LLC for apiary access. As per the manufacturer’s instructions, 0.5 ml of UBeeO was applied to a small circular region of capped, non-emerging honey bee brood cells, and hygienic response was quantified after two hours. Assay scores were calculated as the percentage of the capped cells at T0 that were manipulated (any uncapping including piercing) at T_2_. Colonies that tested over 60% were considered hygienic [58].

### Innate immune-response trials

To evaluate differences in innate immune response, behavior, and mortality between hygienic and non-hygienic bees infected with *N. ceranae*, we selected four hygienic colonies (scoring >60% on UBeeO assay) and four non-hygienic colonies (scoring <60% on UBeeO assay) from which to source newly emerged adults. Ten newly emerged adults per colony were collected from frame cages and tested for *Nosema* spp. using standard microscopy (methods above) to ensure an absence of infection at the start of the experiment. Replicate hoarding cages (30 bees per cage) from each colony were randomly assigned a high dose (5 × 10^4^ spores per 0.04 mL), low dose (10^4^ spores per 0.04 mL), or control (sugar only) inoculant, consumed *ad libitum,* for 24 hours [57]. To determine the amount of sugar syrup inoculant consumed, feeders were weighed before and after they were administered to each hoarding cage. Mortality was recorded for each cage, and four adults were extracted every two days post-inoculation for 10-14 days to assess innate immune response. Samples were stored at -80° C until processed [53].

#### Nosema ceranae, Vitellogenin, and Hymenoptaecin quantification

To quantify active *N. ceranae* infection, and *Vitellogenin* and *Hympenoptaecin* gene expression in bees from our innate immune-response trials, relative qPCR analyses were conducted. RNA was extracted from frozen bees using Trizol (Sigma Aldrich) following manufacturers’ instructions and resuspended in 50 ul molecular-grade water. RNA quantity and quality was assessed with a Nanodrop800 spectrophotometer. DNase treatment with amplification-grade DNAseI (Thermofisher) and cDNA synthesis was performed using Invitrogen SuperScript II Reverse Transcriptase (Thermofisher) with dT and random priming. One μl cDNA was amplified by qPCR using SsoAdvanced Universal SYBR Green Supermix (Biorad) in a 20 ul reaction as per manufacturer’s protocol. Cycling parameters were the same for all targets: 95°C for 3 min, 50 cycles of: 95°C for 5 sec, 60°C for 30 sec, followed by a melt curve to assess product specificity. Primers for all targets are listed in S1 and S2 Tables.

### Data Analysis

Data analysis was conducted in R (version 4.5.2). All mixed models were fit using the LME4 package (v1.1.37;[59]). The significance of main effects and interaction terms was assessed using Type II Wald chi-square tests conducted using the *Anova* function from the CAR package (v3.1.3; [60]). We used *alpha* = 0.05 as the threshold of significance. All outliers were retained in our datasets. Non-normality and zero-inflated data were handled using log transformations or modeling with the appropriate distributions.

Colony hygienic status was determined by converting our continuous variable of UBeeO score (0.00-100%) to a binary variable where colonies were considered ‘hygienic’ if UBeeO score >= 60% and ‘non-hygienic’ if UBeeO score < 60%. Prevalence was calculated from presence–absence data as the proportion of bees testing positive for *N. ceranae* or *Nosema* spp., by dividing the number of infected individuals by the total sample size within each experimental group. Differences in prevalence among groups were evaluated using a chi-square test. The terms “load” or “average load” refer to the quantity of *N. ceranae* spores per bee. The term “levels” refers to relative quantification using ΔΔCt and does not imply an absolute unit of measurement.

Relative *N. ceranae* infection levels and Vg/Hym gene expression were quantified using the ΔΔCt method [61]. Ct values for the target genes and *N. ceranae* were normalized to our reference housekeeping gene (*Actin*) to obtain ΔCt values, then compared to the mean ΔCt of the control group to calculate ΔΔCt (1). Relative *N. ceranae* infection, Vg, and Hym expression levels were log-transformed to address non-normal distributions while preserving true zeros.

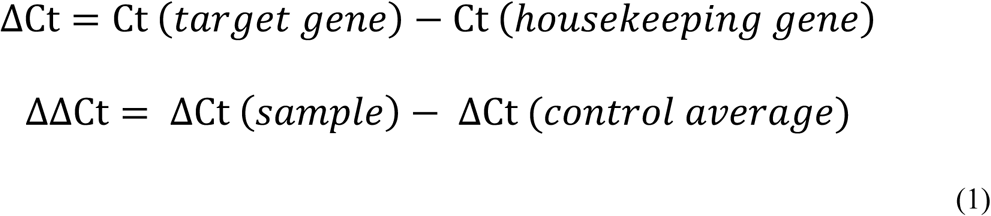

### Inoculant Consumption

To examine the main effect of colony hygienic status, *N. ceranae* dose, and their interaction effect on the amount of sugar syrup inoculant consumed by bees, colony hygienic status, *N. ceranae* dose (control (0 spores), low dose (10^4^ spores per 0.04 mL), high dose (5 × 10^4^ spores per 0.04 mL)), and an interaction term were included as predictor variables in a linear model. An ANOVA Type II test was performed to compute significance of terms in the linear model. Estimated marginal means (EMMs) were calculated for combinations of colony hygienic status and *N. ceranae* dose using the EMMEANS package (v1.11.2.8) and pairwise comparisons of colony hygienic status were performed within each *N. ceranae* dose. To account for the small sample size of hoarding cages in the study, non-parametric Kruskal-Wallis tests were performed to further compute overall significance of colony hygienic status and *N. ceranae* dose as main effects.

#### Nosema ceranae Levels

To test whether relative *N. ceranae* infection levels were influenced by colony hygienic status, *N. ceranae* dose, sampling day, and all possible two-way interactions, we constructed a linear mixed effects model. To account for repeated measures and potential non-independence among individuals reared in the same hoarding cage, cage identity was included as a random effect. Estimated marginal means (EMMs) were calculated for combinations of colony hygienic status, *N. ceranae* dose, and sampling day using the EMMEANS package (v1.11.2.8). Pairwise comparisons among treatment groups were performed using the Tukey method in the MULTCOMP package.

### Immune Gene Expression

To assess whether relative *Vitellogenin* (Vg) or *Hymenoptaecin* (Hym) gene expression were influenced by main effects of colony hygienic status and *N. ceranae*, four additional linear mixed effects models were constructed (two models per target gene). For each target gene, one model included *N. ceranae* dose, colony hygienic status, sampling day, and all possible two-way interaction terms as predictors. Another linear mixed effects model included *N. ceranae* infection levels and colony hygienic status as main effects with an interaction term, and included sampling day as an additive main effect. Cage identity was included as a random effect in all models. Estimated marginal means (EMMs) were calculated for combinations of predictor variables, while estimated marginal trends were used to compute the regression slopes between groups, using the EMMEANS package. Pairwise comparisons among treatment groups were performed using the Tukey method in the MULTCOMP package.

### Survival Probability

To calculate probability of survival, we performed a Kaplan–Meier survival analysis using the SURVIVAL package (v3.8.3). Kaplan–Meier curves were generated to visualize survival probabilities over time for each factor independently. Statistical differences between survival curves were tested using log-rank tests. To evaluate the relative risk of mortality, we fitted a Cox proportional hazards model with colony hygienic status and *N. ceranae* dose as fixed effects, as well as two Cox proportional hazards models with each predictor variable as an independent fixed effect. Survival time in days was used as the response, with individuals surviving the 14-day observation period treated as right-censored.

## Results

### No *N. ceranae* spores found in pupae

Pupal samples were collected from 28 honey bee colonies with confirmed *Nosema* spp. infections in nurse bees ranging in average spore load from 5×10^4^ to 1.4×10⁶ spores per bee. None of the pupal samples collected were found to contain *N. ceranae* spores. These results indicate that the removal of brood through hygienic behavior would have no direct effect on *N. ceranae* load of the colonies in our target population, since *N. ceranae* infection is likely only present in the adult bee castes.

### Group-fed bees experienced higher *N. ceranae* loads

At 10 days post-inoculation, *N. ceranae* prevalence did not differ between group-fed bees (59.3%) and individually-fed bees (43.5%) (*p* = 0.24), nor between group-fed bees with 10 bees (69.6%) or 30 bees (57%) per cage (*p* = 0.38). Of the infected bees, group-feeding resulted in significantly higher average *N. ceranae* loads (*p* < 0.001), but also greater variance (*p* = 0.021) compared to individual-feeding, which can be found in S3 Fig. The number of bees per hoarding cage (10 or 30 bees) had no significant effect on *N. ceranae* loads (*p* = 0.89). Infected group-fed bees experienced an average *N. ceranae* load of 4.28 × 10⁶ ± 2.13 × 10¹³ spores per bee, whereas individually-fed bees experienced an average load of 8.05 × 10⁵ ± 2.61 × 10¹² spores per bee. For subsequent experiments, we selected the group-feeding method with 30 bees per cage to achieve the highest infection levels and obtain a large sample size, despite the higher observed variation in *N. ceranae* loads. We determined that the group-feeding approach better reflected natural *N. ceranae* transmission dynamics in the hive, which are largely driven by a few highly infected individuals [62].

### Immune function and performance differ between hygienic and non-hygienic bees

In *N. ceranae*-inoculated bees, *N. ceranae* infection levels increased over time, demonstrating successful inoculation methods and effective contraction of the pathogen (*p* < 0.001) (Fig 1). At day 12 post-inoculation, we found a *N. ceranae* prevalence of 50% in bees that received the low *N. ceranae* dose and 81.5% in bees that received the high *N. ceranae* dose. All *N. ceranae-*inoculated bees experienced significantly higher infection levels than control bees (*p* < 0.001), but there was only a marginal difference between the low (10^4^ spores per bee) and high *N. ceranae* (5 × 10^4^ spores per bee) dose groups (*p* = 0.052) (Fig 2). We attribute the low levels of *N. ceranae* observed in 17.2% of control bees at day 12 post-inoculation to newly emerged adults ingesting spores on their original frames before being transferred to hoarding cages for the experiment.

**Fig 1.**
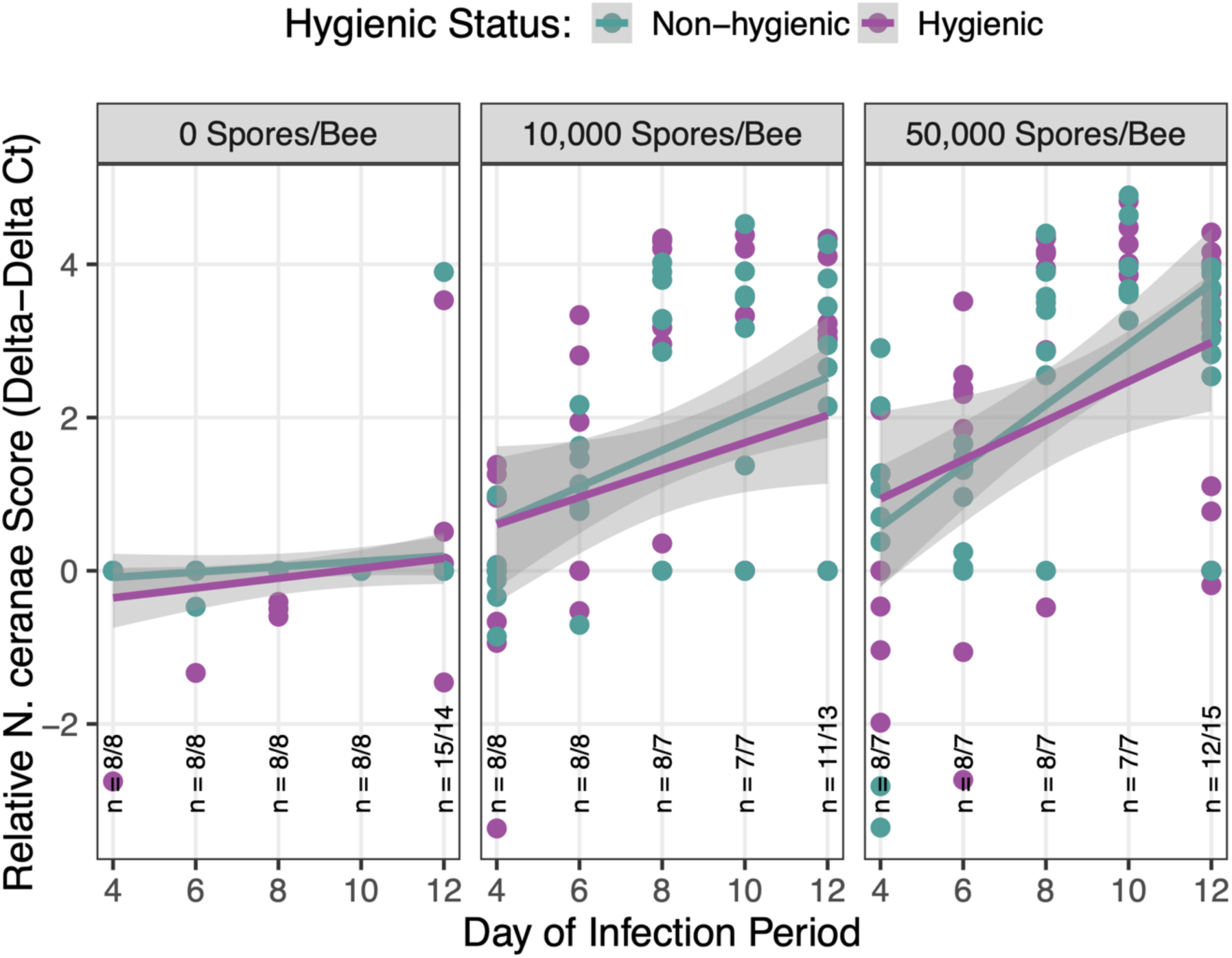
Effect of colony hygienic status and *N. ceranae* dose on relative *N. ceranae* infection levels in bees over time. There was a significant effect of sampling day (*χ^2^*_1_ = 42.87, *p* < 0.001) and *N. ceranae* dose (*χ^2^*_2_ = 79.11, *p* < 0.001) on relative *N. ceranae* infection levels (ΔΔCt, log-transformed), with a significant interaction effect of sampling day and *N. ceranae* dose (*χ^2^*_2_ = 15.56, *p* < 0.001). There was no effect of colony hygienic status (*χ^2^*_1_ = 1.22, *p* = 0.27). Purple points/lines represent hygienic bees; green points/lines represent non-hygienic bees. Panels correspond to *N. ceranae* doses (High = 5 × 10^4^ spores/bee; Low = 1 × 10^4^ spores/bee; Control = 0 spores/bee). Sample sizes are denoted at each time point as *n* = hygienic bees / non-hygienic bees.

**Fig 2.**
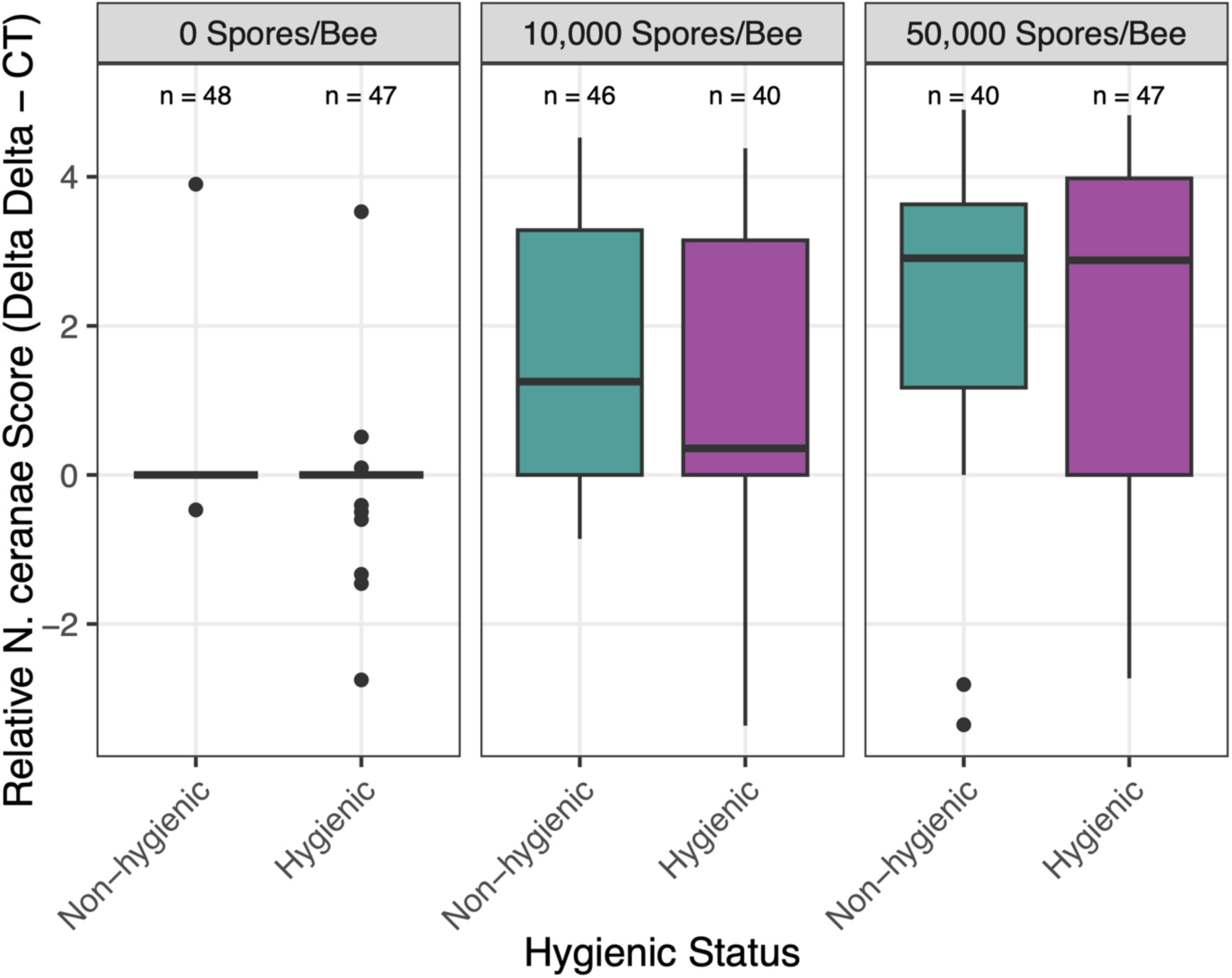
Effect of *N. ceranae* dose on relative *N. ceranae* infection levels in hygienic and non-hygienic bees. There was a significant main effect of *N. ceranae* dose on relative infection levels (*χ^2^*_2_ = 79.11, *p* < 0.001), but no significant effect of colony hygienic status (*χ^2^*_1_ = 1.22, *p* = 0.27) or an interaction effect between colony hygienic status and *N. ceranae* dose (*χ^2^*_2_ = 0.062, *p* = 0.938). Purple boxes represent hygienic bees; green boxes represent non-hygienic bees. Panels correspond to *N. ceranae* doses (High = 5 × 10^4^ spores/bee; Low = 10^4^ spores/bee; Control = 0 spores/bee). Infection levels are shown as relative *N. ceranae* score (ΔΔCt, log-transformed). Sample sizes are denoted above each box and varied based on bee mortality. Significance between pairs is denoted as ‘.’ < 0.1, ‘*’ < 0.05, ‘**’ < 0.01, ‘***’ < 0.001.

Hygienic and non-hygienic bees did not differ significantly by *N. ceranae* prevalence (*p* = 1) nor in increasing *N. ceranae* infection levels observed over time (*p* =0.938). However, we found that hygienic bees in hoarding cages consumed significantly less sugar syrup inoculant in the 24hr inoculation period overall, compared to non-hygienic bees (*p* < 0.001). There was no significant effect of *N. ceranae* dose (*p* = 0.129) nor an interaction effect between colony hygienic status and *N. ceranae* dose (*p* = 0.101). Broken out by *N. ceranae* dose in a pairwise comparison of our linear model, we found a biologically relevant trend that hygienic bees consumed less of the highest *N. ceranae* dose (5 × 10^4^ spores per bee) compared to non-hygienic bees (*p* = 0.055), but otherwise no significant differences were found between the volume of sugar syrup consumed by hygienic and non-hygienic bees among the control (0 spores per bee, *p* = 0.595) and low *N. ceranae* dose groups (1 × 10^4^ spores/bee, *p* = 0.449) (Fig 3).

**Fig 3.**
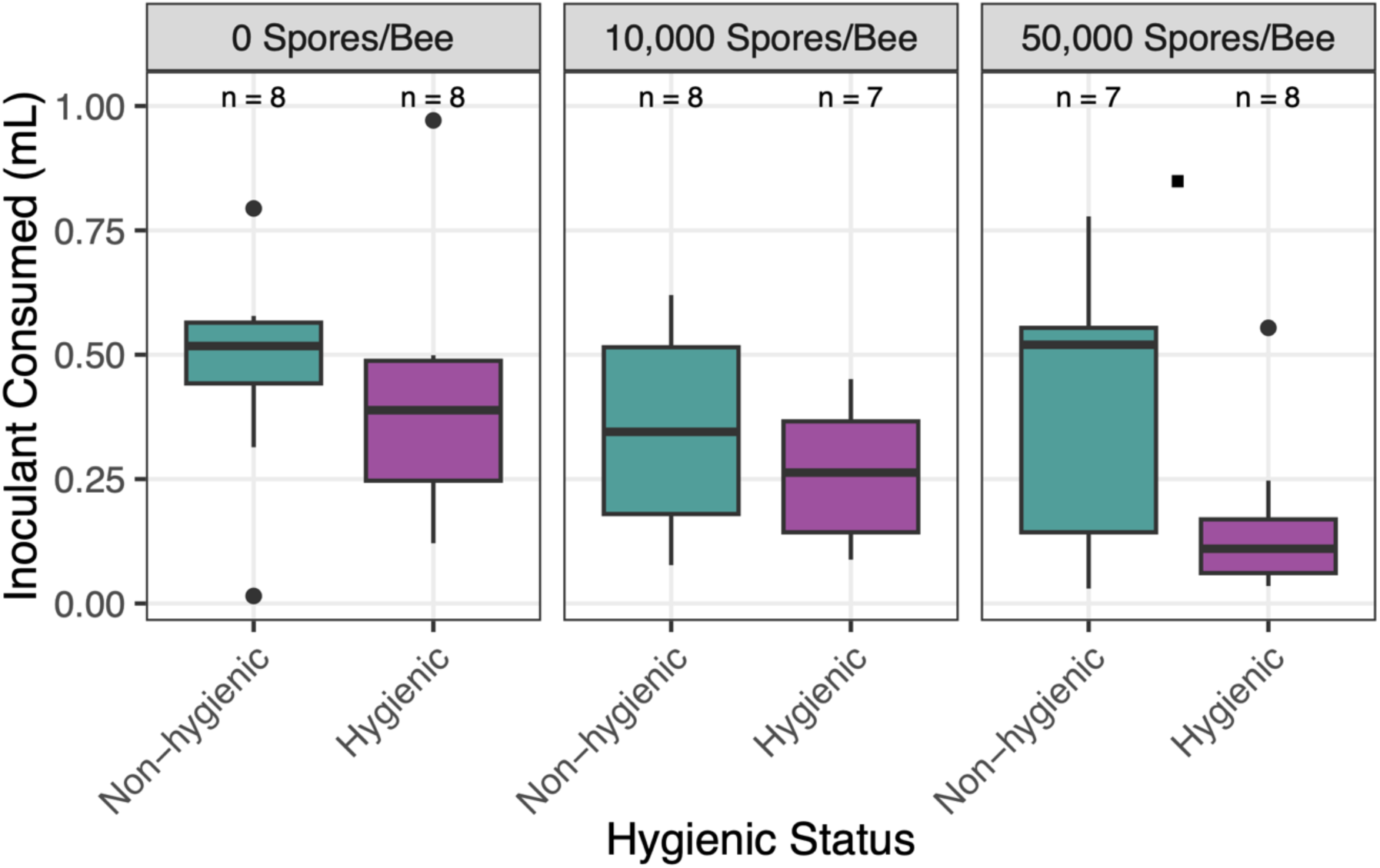
Effect of colony hygienic status and *N. ceranae* dose on the volume of sugar syrup inoculant consumed by bees in hoarding cages. There was a significant main effect of colony hygienic status on the volume of inoculant consumed by bees in hoarding cages (Kruskal-Wallis test, χ^2^ = 4.3104, *df* = 1, *p* = 0.038), but no effect of *N. ceranae* dose (χ^2^ = 4.0909, *df* = 2, *p* = 0.129) nor interaction between colony hygienic status and *N. ceranae* dose (χ^2^ = 9.228, *df* = 5, *p* = 0.101). There was a marginal difference in the volume of inoculant consumed between hygienic and non-hygienic bees in the high *N. ceranae* dose group (*p* = 0.055). Purple boxes represent hygienic bees; green boxes represent non-hygienic bees. Panels correspond to *N. ceranae* doses (High = 5 × 10^4^ spores/bee; Low = 1 × 10^4^ spores/bee; Control = 0 spores/bee). Inoculant consumption is shown in milliliters (mL). Sample sizes are denoted above each box and varied based on bee mortality. Significance between pairs is denoted as ‘.’ < 0.1, ‘*’ < 0.05, ‘**’ < 0.01, ‘***’ < 0.001.

*Vitellogenin* expression levels significantly differed by colony hygienic status (*p* = 0.006) and sampling day (*p* < 0.001) (Fig 4). At peak *N. ceranae* infection (day 12 post-inoculation), hygienic bees showed upregulated *Vitellogenin* (Vg) expression while non-hygienic bees showed downregulated expression in all groups, including the low *N. ceranae* dose (*p* < 0.001), high *N. ceranae* dose (*p* = 0.005), and control (*p* < 0.001) bees. Additionally, we found a marginal difference in Vg expression on day 4 post-inoculation between hygienic and non-hygienic bees that received the highest dose of *N. ceranae* (5 × 10^4^ spores per bee) (*p* = 0.056). Conversely, we found that *Hymenoptaecin* levels did not differ significantly by colony hygienic status, *N. ceranae* dose, sampling day, nor their interactions, which can be found in S4 Figure.

**Fig 4.**
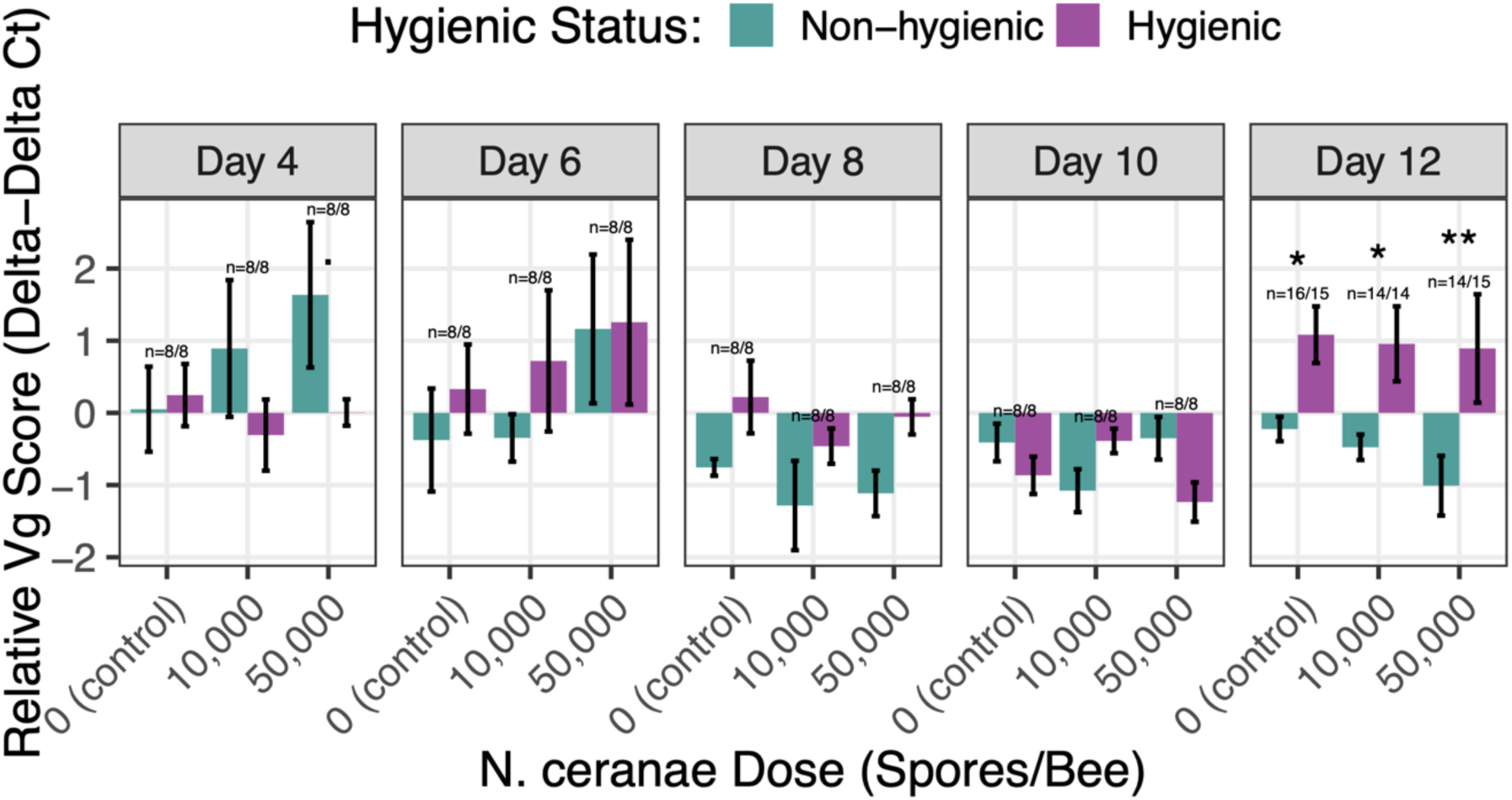
Effect of colony hygienic status on *Vitellogenin* (Vg) gene expression across time and *N. ceranae* doses. There were significant main effects of colony hygienic status (χ^2^_1_ = 7.62, *p* = 0.006) and sampling day (χ^2^_4_ = 22.78, *p* < 0.001), as well as a significant interaction between colony hygienic status and day (χ^2^_4_ = 19.79, *p* < 0.001) on *Vitellogenin* expression levels. *Vitellogenin* expression levels are shown as the relative *Vg* score (ΔΔCt, log_1_₀-transformed). Purple bars indicate hygienic bees; green lines indicate non-hygienic bees. Panels correspond to sampling days post-inoculation. Error bars represent standard errors of the mean. Sample sizes are denoted above each bar pair as *n* = hygienic bees/non-hygienic bees. Significance between pairs is denoted as ‘.’ < 0.1, ‘*’ < 0.05, ‘**’ < 0.01, ‘***’ < 0.001.

As a more accurate measure of bees’ immune response against *N. ceranae*, we correlated *Vitellogenin* (Vg)*/Hymenoptaecin* (Hym) expression levels with actual *N. ceranae* infection. We found that *N. ceranae* infection levels (*p* = 0.028) and colony hygienic status (*p* = 0.012) had a significant main effect on Vg expression levels, but no significant interaction effect (*p* = 0.242). In all bees, Vg expression levels downregulated in response to increasing *N. ceranae* infection levels; however, hygienic bees showed less of a negative relationship, where Vg expression was relatively unaffected by increasing *N. ceranae* infection levels (Fig 5). Evaluating *Hymenoptaecin* expression in response to *N. ceranae* infection, we found a significant interaction effect between *N. ceranae* infection levels and colony hygienic status (*p* = 0.016), where hygienic bees showed lower Hym expression with mild infection followed by upregulation in response to increasing *N. ceranae* infection levels. In contrast, non-hygienic bees showed consistent downregulation in Hym expression in response to *N. ceranae* infection levels (Fig 6).

**Fig 5.**
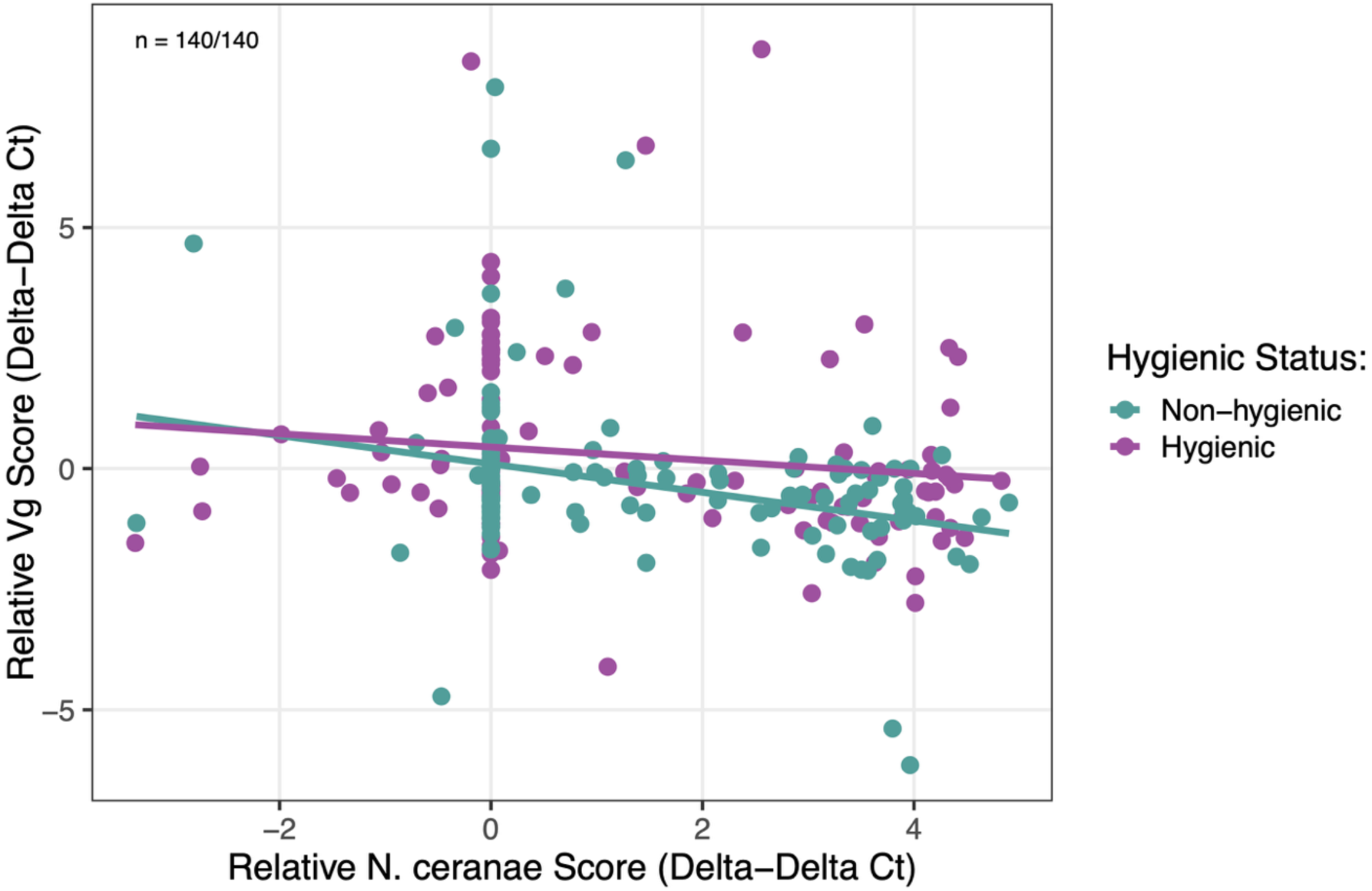
Effect of *N. ceranae* infection levels on *Vitellogenin* (Vg) gene expression by colony hygienic status. *Vg* gene expression levels (ΔΔCt, log-transformed) were impacted significantly by main effects of colony hygienic status (χ^2^_1_ = 6.27, *p* = 0.012), *N. ceranae* infection levels (χ^2^_1_ = 4.81, *p* = 0.028), and sampling day (χ^2^_4_ = 14.27, *p* = 0.006). There was no significant interaction effect between colony hygienic status and *N. ceranae* infection levels (χ^2^_1_ = 1.37, *p* = 0.242). Purple points/lines represent hygienic bees; green points/lines represent non-hygienic bees. Sample sizes are denoted as *n* = hygienic bees / non-hygienic bees.

**Fig 6.**
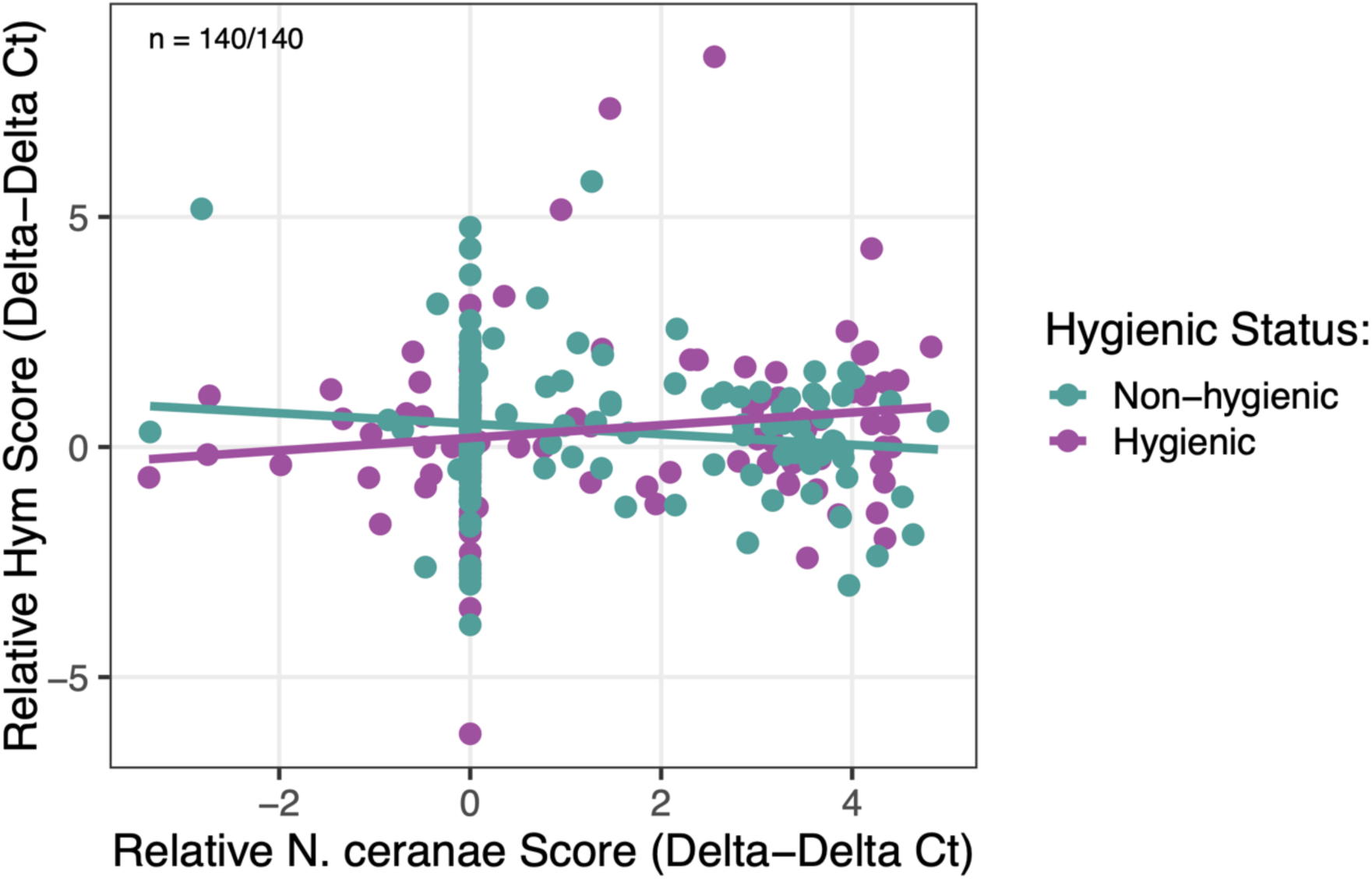
Effect of *N. ceranae* infection levels on *Hymenoptaecin* (Hym) gene expression by colony hygienic status. *Hym* gene expression levels (ΔΔCt, log_1_₀-transformed) were impacted by an interaction effect of colony hygienic status and *N. ceranae* infection levels (χ^2^_1_ = 5.805, *p* = 0.016). No main effects of *N. ceranae* infection levels (χ^2^_1_ = 0.05, *p* = 0.831) or colony hygienic status (χ^2^_1_ = 0.00, *p* = 0.997) on Hym expression levels were detected. Purple points/lines represent hygienic bees; green points/lines represent non-hygienic bees. Sample sizes are denoted as *n* = hygienic bees / non-hygienic bees.

Hygienic and non-hygienic bees differed significantly in their probability of survival during the experimental trials (*p* = 0.02). Among the *N. ceranae*-inoculated groups, hygienic bees had significantly better survival odds than non-hygienic bees starting 8 days post-inoculation (*p* = 0.004), where non-hygienic bees had a 53% higher risk of death when infected (Fig 7). Bees in the control group did not differ in survival probability based on colony hygienic status (*p* = 0.50). Among all bees, *N. ceranae* inoculation significantly affected the probability of bee survival (*p* < 0.001) starting four days post-inoculation. *Nosema ceranae*-inoculated bees had a 135-138% higher risk of death compared to control bees, but the high and low *N. ceranae* dose groups did not differ significantly in survival probability (*p* = 0.93) (Fig 8).

**Fig 7.**
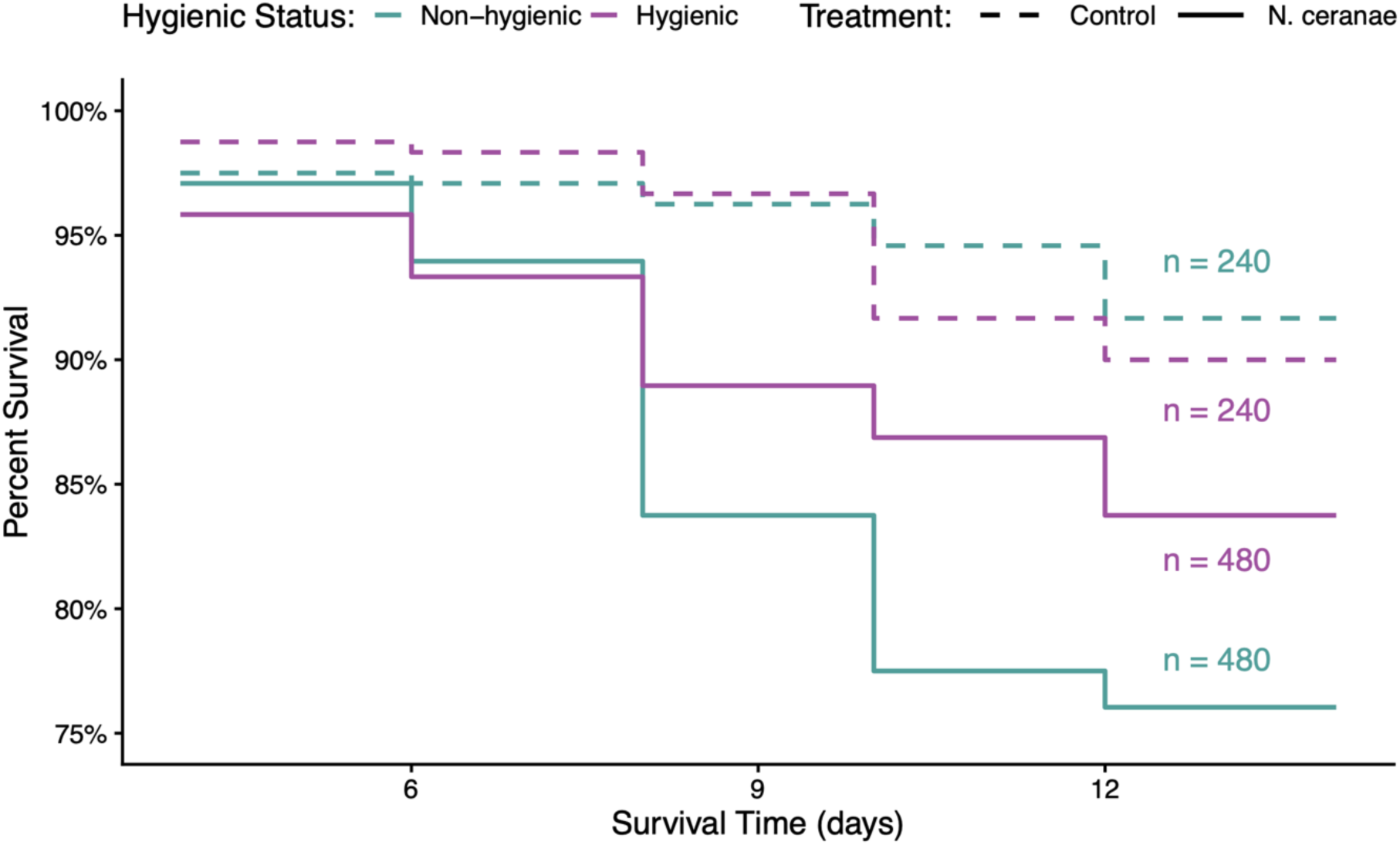
Survival probability of bees by colony hygienic status and treatment group over time. Among *N. ceranae*-inoculated groups, hygienic bees had significantly better survival odds than non-hygienic bees (χ^2^ = 8.3, df = 1, *p* = 0.004; HR = 1.53, 95% CI: 1.15–2.04). Among control bees, there was no significant difference by colony hygienic status (χ^2^ = 0.4, df = 1, *p* = 0.50; HR = 0.83, 95% CI: 0.46–1.50). Purple lines indicate hygienic bees; green lines indicate non-hygienic bees. Dashed lines represent control bees; solid lines represent *N. ceranae*-inoculated bees. Sample sizes are denoted beside each line.

**Fig 8.**
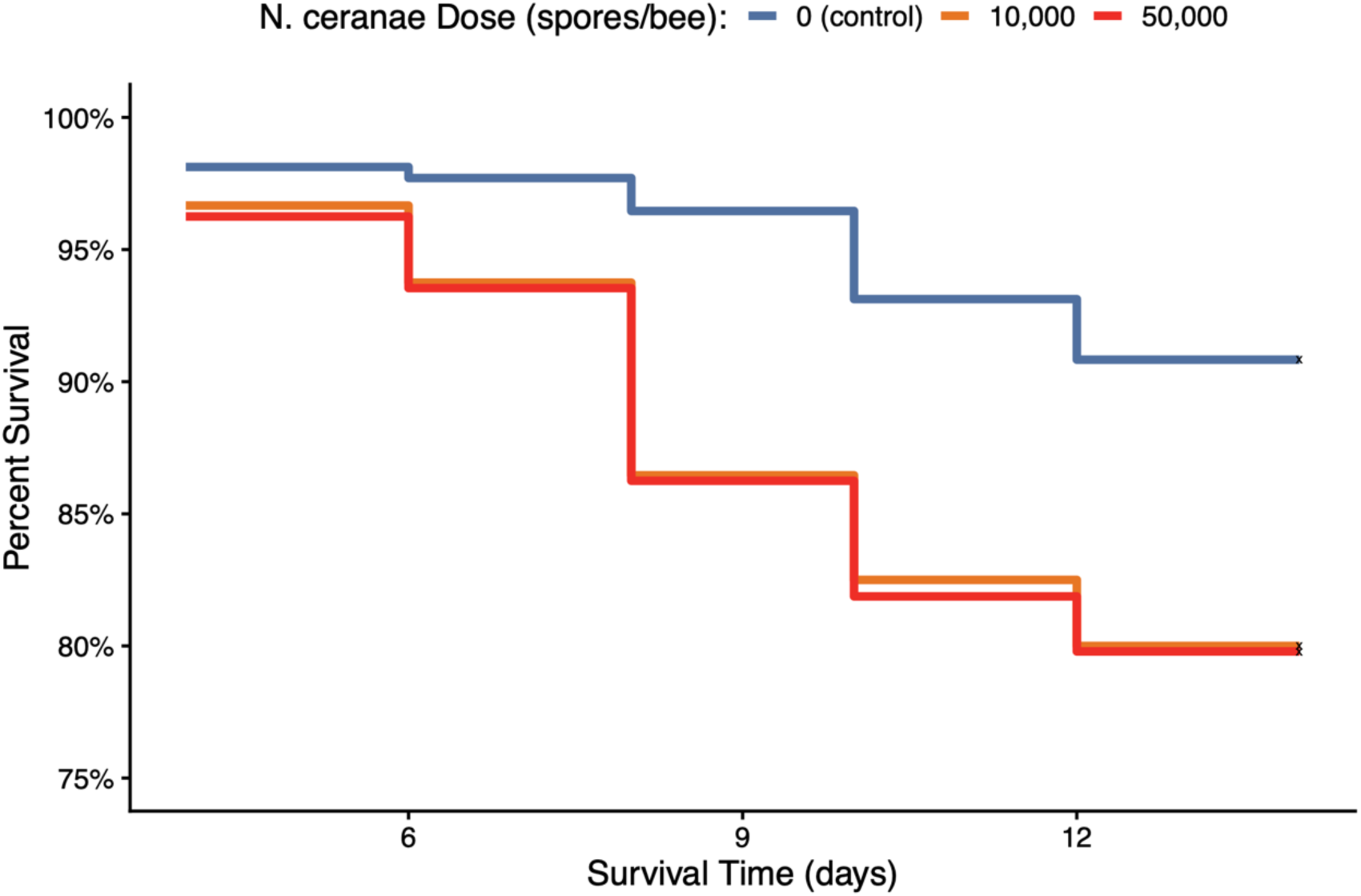
Effect of *N. ceranae* dose on survival probability over time. There was a significant effect of *N. ceranae* dose on survival probability (χ^2^_2_ = 28.3, df = 2, *p* = 7 × 10⁻⁷), where *N. ceranae* exposure significantly increased the hazard of death compared to control bees. Bees that received the low and high dose *N. ceranae* inoculant did not differ significantly in survival probability (*p* = 0.93). Red line indicates high *N. ceranae* dose (5 × 10^4^ spores/bee), orange line indicates low *N. ceranae* dose (10^4^ spores/bee), and blue line indicates control group (0 spores/bee). Sample sizes are *n* = 240 for each group.

## Discussion

Our results suggest that individual bees originating from hygienic (high UBeeO) colonies express innate defense mechanisms against *Nosema ceranae*. Despite hygienic bees showing similar levels of *N. ceranae* infection to non-hygienic bees in this cage study, we find that individual bees in hygienic colonies may actively mitigate *N. ceranae* infection by 1) limiting the amount of inoculant consumed, 2) upregulating *Vitellogenin* expression during peak infection, 3) upregulating *Hymenoptaecin* expression in response to increasing infection, and 4) experiencing greater survivability. Hygienic bees may also modulate investment in innate immunity in response to infection severity while limiting the health impacts of *N. ceranae* by maintaining Vg and Hym expression as infection increases. It is important to note that, due to our study design, bees were limited in their ability to remove unhealthy individuals from their cages—a key behavior that likely contributes to reducing *N. ceranae* loads in hygienic colonies under natural hive conditions. As a result, our measurements of individual *N. ceranae* levels may not fully reflect differences between hygienic and non-hygienic colonies in the field since social immunity is known to play a significant role in host-pathogen dynamics.

We found no evidence of *N. ceranae* spores in developing pupae of *N. ceranae-*infected colonies, suggesting that brood does not likely experience *N. ceranae* infection under natural hive settings in our target population. Therefore, we believe that hygienic behavior would have no direct effect on the level of *N. ceranae* infection in the colony, since the behavior acts solely on infected brood. Previous studies have shown that larvae can be manually inoculated with *N. ceranae* and will develop *N. ceranae*-induced physiological and metabolic impairments as adults [34,35]; however, pupae have only been shown to experience rare infection under natural hive conditions [36,37]. Overall, *Nosema ceranae* infections in brood have not been thoroughly investigated, and the extent of their prevalence—and how it may vary geographically—remains unclear. Our findings are consistent with previous studies showing an absence of *N. ceranae* infection in newly emerged adults [17,53] and little direct impact of hygienic behavior on *N. ceranae*, besides an observed overall reduction at the apiary level over time [14,40].

Compared to non-hygienic bees, hygienic bees consumed less sugar syrup inoculant overall. We found a biologically relevant trend that hygienic bees consumed less of the high dose *N. ceranae* inoculant, compared to the low *N. ceranae* dose or control group. It remains unclear whether hygienic colonies may be able to detect and avoid *N. ceranae-*contaminated food sources within the hive. *N. ceranae* transmission most often occurs through the oral-fecal route, by consuming contaminated food stores [63,64], cleaning bee excrement from the frames, or through engagement in trophallaxis with infected nestmates [65]. The recognition and avoidance of *N. ceranae* spores on hive materials could have a significant impact on reducing *N. ceranae* transmission in the colony. Our findings point to a potentially heightened sensitivity of hygienic bees to atypical odors at high concentrations, such as those associated with *N. ceranae* or other pathogens. Future studies should investigate whether hygienic bees avoid *N. ceranae*–contaminated hive materials in cage-choice experiments and how they respond to pathogen-related odors in olfactometer assays. Observational studies should also assess whether hygienic bees exhibit additional in-hive behaviors, such as the “entombment” of contaminated food stores [66] increased attentiveness to infected adults, or higher rates of auto- and allo-grooming [1,6].

Overall, *N. ceranae* infection substantially increased the risk of bee mortality, supporting existing evidence of its negative impact to bee health [19–22,25,27]. However, hygienic bees seem to be more tolerant to *N. ceranae* compared to non-hygienic bees. Despite exhibiting similar *N. ceranae* infection levels in our cage study, hygienic bees survived significantly longer than non-hygienic bees when infected with *N. ceranae*. Tolerance is defined by an organism’s ability to minimize the damage caused by a pathogen, rather that reducing or eliminating the pathogen itself. Our findings are consistent with the enhanced survival observed in infected drones of a known *N. ceranae*-tolerant honey bee strain in Denmark [26,67]. When infected with *N. ceranae*, tolerant drones showed normal rates of apoptosis in the midgut epithelium, maintaining normal cell function and the ability to rid damaged tissue. Limiting *N. ceranae’s* ability to inhibit apoptosis– a key mechanism in the pathogenesis of *N. ceranae* infections– may therefore contribute to the enhanced survivability observed in hygienic bees. If an altered apoptotic response to *N. ceranae* infection in hygienic bees could facilitate defecation of infected cells outside of the hive, it may also play a role in limiting transmission of the pathogen between nestmates and explain the reduced loads observed at the colony level [3]. However, further evaluations to compare the apoptosis rates between hygienic and non-hygienic bees with *N. ceranae* infections are needed to support these hypotheses.

Improved survivability in hygienic bees may also be explained by an enhanced buffering capacity that reduces the energetic stress caused by *N. ceranae* [19,20], as demonstrated in *N. ceranae*-tolerant drones in Denmark [3]. Although we did not measure sugar syrup consumption throughout the 12-day infection period, the reduced overall intake of inoculant during the inoculation period may indicate a lower carbohydrate demand in hygienic bees. Future studies should evaluate hemolymph trehalose levels in *Nosema*-infected hygienic bees to better understand if their energy availability is preserved over time [68], which may contribute to their improved survival and performance in the colony. Nevertheless, the prolonged survival of *N. ceranae-*infected individuals in hygienic colonies may help to alleviate colony-level impacts of *N. ceranae* (e.g. reduction in population, decreased honey production [15]) by retaining the workforce over time. Conversely, surviving beyond seven days old, when precocious foraging caused by *N. ceranae* infection is likely to occur [24], may be an adaptation of hygienic colonies to lower pathogen transmission in the hive by favoring the reduced homing ability of infected adults [23,69].

We evaluated *Vitellogenin* and *Hymenoptaecin* gene expression to assess the innate immune function of hygienic bees with *N. ceranae* infection. Hygienic bees exhibited significantly upregulated *Vitellogenin* (Vg) gene expression at peak *N. ceranae* infection (day 12 post-inoculation). Notably, the level of upregulation was independent of *N. ceranae* dose, indicating that hygienic bees exhibit upregulated Vg expression at this time point even in the absence of pathogen exposure. Non-hygienic bees showed downregulated Vg expression as *N. ceranae* levels increased compared to hygienic bees, which maintained relatively constant Vg expression, suggesting that non-hygienic bees may have a more compromised or altered physiological response under infection compared to hygienic bees. The pronounced increase in Vg expression observed at day 12 in hygienic bees has not been reported in previous studies. In a healthy colony, Vg levels typically peak in nurse bees around four days old and decline as workers transition from in-hive tasks to foraging duties [70,71]. While *N. ceranae* infection can cause elevated Vg levels in younger bees, the late spike observed in hygienic bees is unusual and suggests potential functional implications that warrant further investigation.

The overexpression of Vg at day 12 post-inoculation (≈15 days old) could reflect changes in normal behavioral maturation [50] and social organization [51] in hygienic colonies. However, overexpression could also enhance immunological defenses and resilience against pathogens. *Vitellogenin* has been shown to bind to pathogens, suppress microbial growth, and contribute to tissue repair from oxidative stress [48]. At 15 days old, workers typically transition to undertaker roles [72], or in hygienic colonies, perform hygienic behaviors to remove dead or parasitized individuals [73]. Concurrent Vg upregulation may protect these bees while performing risky duties. High levels of Vg are linked to the prolonged lifespan of queen bees and stress resilient *diutinus* bees [52], suggesting that upregulated Vg may underlie the enhanced survivability observed in hygienic bees. Further, the potential for Vg to perform trans-generational immune priming functions [49,74] could have important implications for the heritability of *N. ceranae* tolerance in hygienic colonies.

While hygienic bees do not show clear differences in *Hymenoptaecin* expression when compared to non-hygienic bees over time, there is a significant relationship between Hym expression levels and *N. ceranae* infection severity. Hygienic bees exhibited lower Hym expression at low infection intensities compared to non-hygienic bees and a stronger upregulation of Hym expression in response to increasing *N. ceranae* infection levels. Our findings suggest that hygienic bees may reduce investment in innate immunity under low pathogen stress, but are better able to combat *N. ceranae* as infection increases. In contrast, non-hygienic bees show a stronger immune response under low pathogen stress but weaken in Hym expression as *N. ceranae* infection increases. The relationship between Hym expression and *N. ceranae* infection in non-hygienic bees reflects previous studies showing the pathogen’s immunosuppressing capabilities in infected bees [22,53]. While we do not find higher *Hymenoptaecin* expression in hygienic bees overall, our findings are comparable with previous work showing elevated *Toll* pathway–mediated immune gene expression in *N. ceranae*–tolerant drones [67] and higher Hym expression in workers from hygienic colonies [12]. Furthermore, the reduced initial investment in Hym expression in hygienic bees may result in more energy availability and explain their reduced demand for sugar syrup inoculant during the inoculation period. Future studies should examine additional *Toll* pathway–mediated antimicrobial peptides to fully characterize innate immunity in hygienic bees and its role in controlling *N. ceranae*.

Our study highlights that colony-level resistance to *N. ceranae* may emerge from the cumulative effects of individual-level mechanisms. Disease resistance that confers reduced levels of pest and pathogen infestation at the colony level has been a major focus in recent honey bee breeding efforts [75–77]. However, selective breeding programs may also consider targeting individual-level tolerance mechanisms, such as maintained apoptosis rates and/or improved energetic buffering capacity under infection, that avoid antagonistic host–parasite coevolution [78] and could promote colony-level resistance to pathogens while circumventing the pitfalls of pure tolerance-based selection [79]. For example, individual tolerance to Deformed Wing Virus is thought to contribute to the winter survival of *Varroa*-resistant colonies [80]. In general, tolerance mechanisms are not usually pathogen-specific and could offer protection against a broad range of pathogens in honey bee colonies [77].

Overall, our findings suggest that hygienic behavior in honey bee colonies, quantified by the UBeeO assay, may be linked to individual-level defenses that function concurrently to maintain low levels of *N. ceranae* at the colony level. These investigations advance our understanding of how hygienic behavior performance can predict pathogen loads and have important implications for selective breeding strategies, *N. ceranae* prevention, and disease management. Further research is needed to explore potential social immune mechanisms that combat *N. ceranae* and how nestmates interact with infected individuals in hygienic colonies. While previous studies have shown that nestmates can exhibit behaviors ranging from avoidance to aggression towards *Nosema*-infected individuals [81], it remains unclear how these interactions differ in hygienic colonies and how social behaviors might complement the individual-level traits of hygienic bees revealed in this study. Here, we offer a valuable perspective on the abilities of individual workers to regulate *N. ceranae* infection in hygienic colonies and contribute to ongoing efforts to improve honey bee health.

## Acknowledgments

We thank the USDA Beltsville technicians, Allison Shaulis and Kyle Grubbs, for their assistance with laboratory work. We are especially grateful to Michael Palmer of French Hill Apiaries and Bianca Braman of Vermont Bees LLC for providing access to their apiaries and extensive support throughout the project.

## Supporting Information

**S1 Table.**
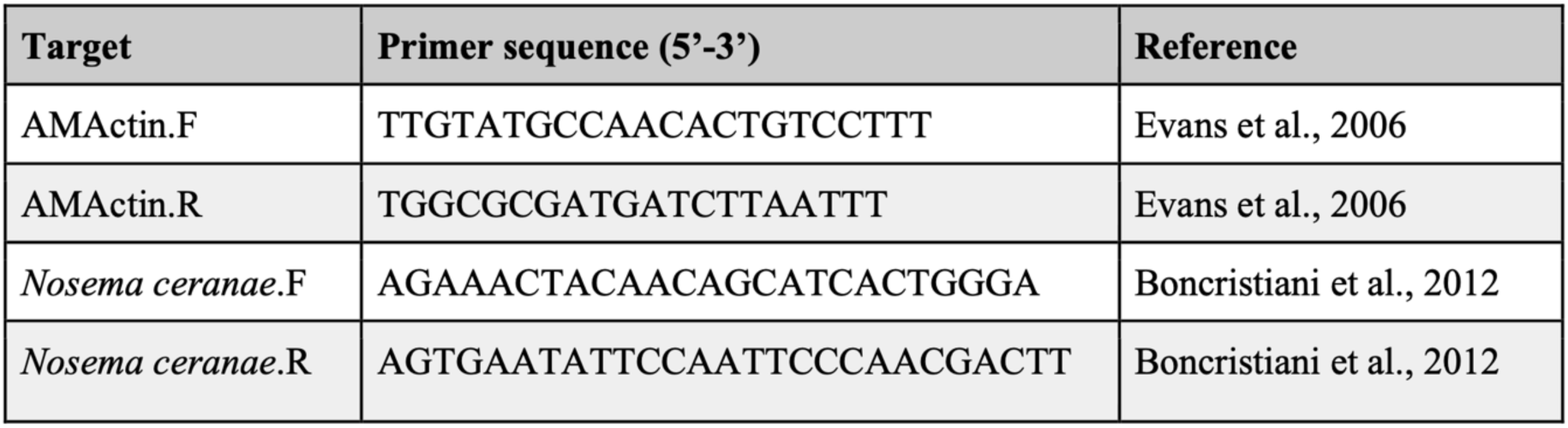
Primers used to determine relative quantification of *Nosema ceranae* [82].

**S2 Table.**
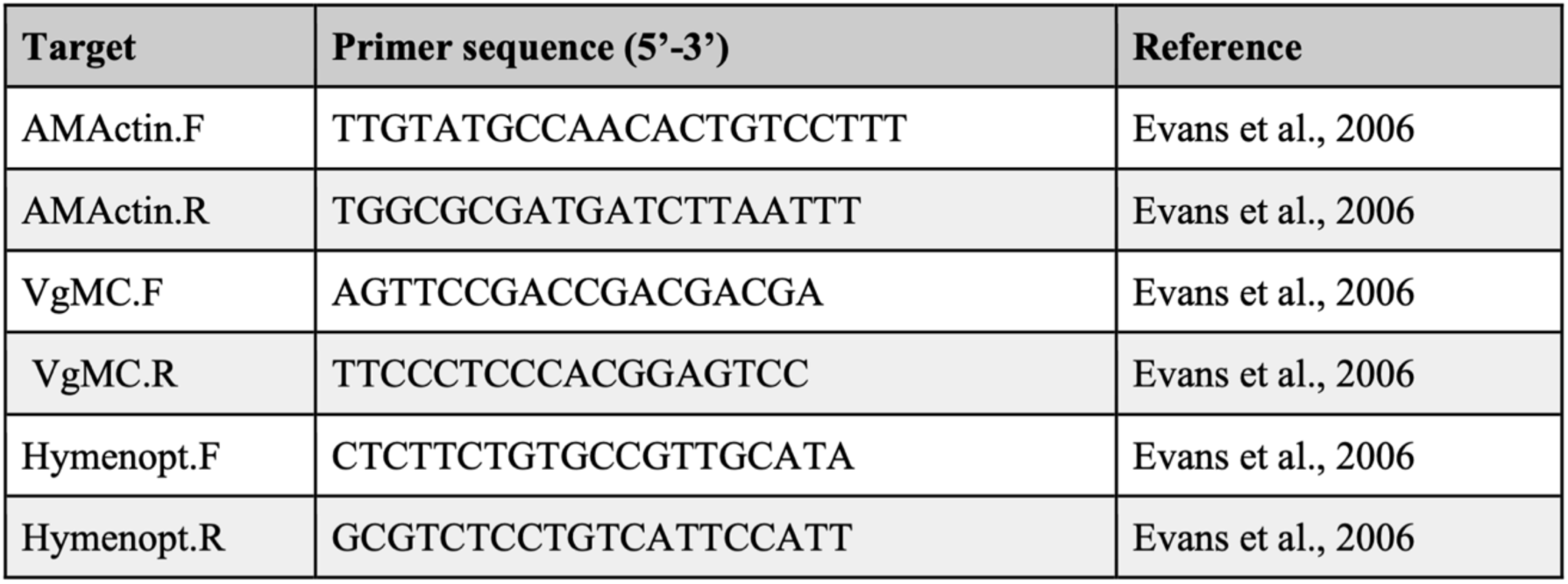
Primers used to determine relative quantification of *Hymenoptaecin* and *Vitellogenin* expression [1].

**S3 Fig.**
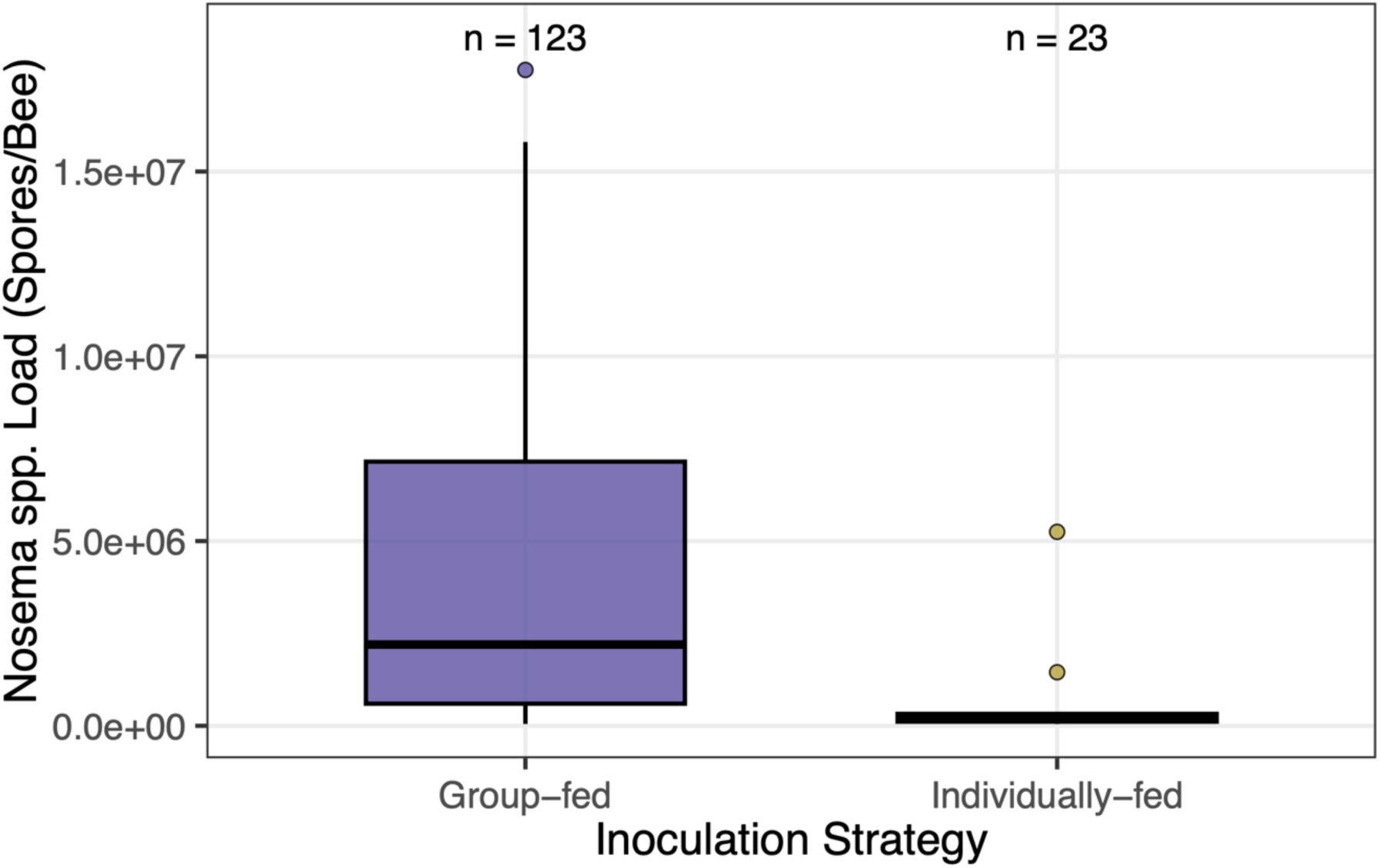
Effect of inoculation strategy on *Nosema* spp. loads in worker bees. *Nosema* spp. loads (spores per bee) differed significantly between group-fed and individually fed bees (Welch’s t-test: t₃₅.₀₅ = 4.67, p < 0.001), with greater loads and variance among group-fed bees (Levene’s test: F_1_,₈_1_ = 5.52, p = 0.021). Purple boxes indicate group-fed bees, and yellow boxes indicate individually-fed bees. Sample sizes are denoted above each box.

**S4 Fig.**
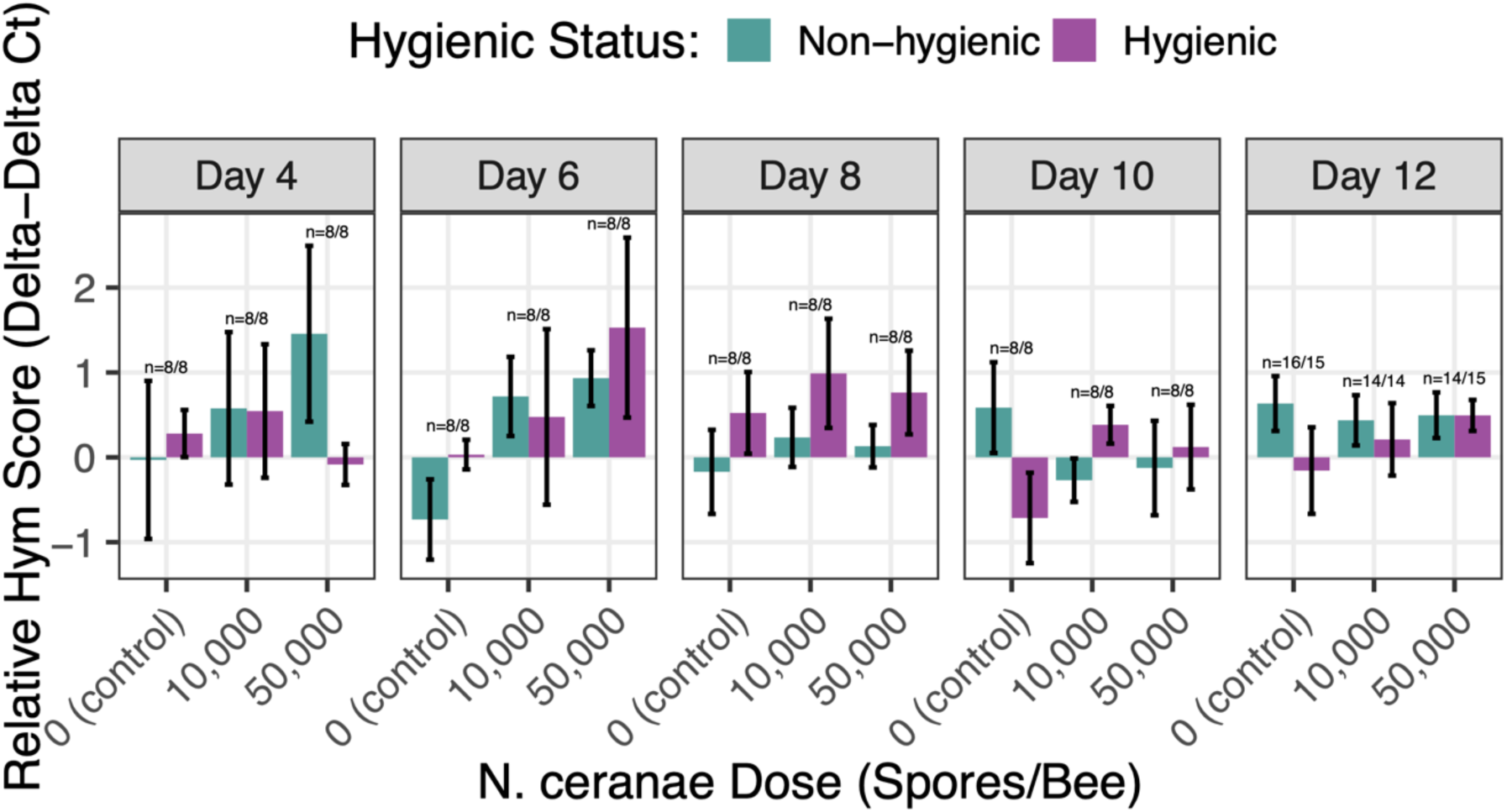
Effect of colony hygienic status on *Hymenoptaecin* (Hym) gene expression across time and *N. ceranae* doses. No significant main effects of colony hygienic status (*χ^2^*_1_ = 0.012, *p* = 0.913), sampling day (*χ^2^*_4_ = 3.33, *p* = 0.504), *N. ceranae* dose (*χ^2^*_2_ = 3.23, *p* = 0.199), nor interaction effects between the predictor variables were detected. *Hymenoptaecin* expression levels are shown as the relative *Hym* score (ΔΔCt, log_1_₀-transformed). Purple bars represent hygienic bees; green bars represent non-hygienic bees. Error bars represent standard errors of the mean. Sample sizes are denoted above each bar pair as *n* = hygienic bees/non-hygienic bees. Significance between pairs is denoted as ‘.’ < 0.1, ‘*’ < 0.05, ‘**’ < 0.01, ‘***’ < 0.001.

